# Transport among protocells via tunneling nanotubes

**DOI:** 10.1101/2021.09.16.460285

**Authors:** Ingrid Jin Schanke, Lin Xue, Karolina Spustova, Irep Gözen

**Affiliations:** Centre for Molecular Medicine Norway, Faculty of Medicine, University of Oslo, 0318 Oslo, Norway; Department of Chemistry, Faculty of Mathematics and Natural Sciences, University of Oslo, 0315 Oslo, Norway

**Keywords:** protocell, lipid nanotube, molecular transport, origin of life

## Abstract

We employ model protocell networks for evaluation of molecular transport through lipid nanotubes as potential means of communication among primitive cells on the early Earth. Network formation is initiated by deposition of multilamellar lipid reservoirs onto a silicon oxide surface in an aqueous environment. These reservoirs autonomously develop into surface-adhered protocells interconnected via lipid nanotubes, and encapsulate solutes from the ambient buffer in the process. We prepare networks in the presence of DNA and RNA and observe encapsulation of these molecules, and their diffusive transport between the lipid compartments via the interconnecting nanotubes. By means of an analytical model we determine key physical parameters affecting the transport, such as nanotube diameter and compartment size. We conclude that nanotube-mediated transport in self-organized nanotube-vesicle networks could have been a possible pathway of chemical communication between primitive, self-assembled protocells under early earth conditions, circumventing the necessity for crossing the membrane barrier. We suggest this transport within a closed protocell network as a feasible means of RNA and DNA exchange under primitive prebiotic conditions, possibly facilitating early replication.

## Introduction

How the first living cell emerged from prebiotic matter on the early Earth is still an unsolved question. Current studies focusing on this problem utilize synthetic model precursors of primitive cells, the ‘protocells’. Protocells carry features of living cells, but are structurally and functionally much simpler^1^. A feature in common with a modern cell, which is surrounded by a plasma membrane is a biosurfactant bilayer, which establishes a boundary, an identity, and an interface suitable for chemical exchange^2, 3^. A bilayer envelope satisfies one of the three conditions of the Chemoton model, the hypothetical chemical entity which features all necessary criteria of ‘living’^4^. A membranous protocell is typically prepared under laboratory conditions via self-assembly of bulk amphiphiles in an aqueous solution^2, 3^.

In order to cross the boundary between the non-living and living matter, a primitive cell needs to be able to develop, grow and eventually self-replicate, resulting in formation of genetically identical daughter cells. To undergo Darwinian evolution during this process, protocells should attain the ability to sense, and adapt to, relevant changes in the environment. Such changes can include perturbations caused by other protocells. Modern cells, for example, can communicate by secreting chemical signals into their surroundings, which is recognized by nearby cells, that are either in direct contact, or a short distance away^5-9^. The response can be manifold, for example tuning the gene expression to change density of the cell population, or induce cell division. Whether bacteria or eukaryotes, cellular communication and division pathways require the coordination of multiple sets of proteins and ligands^7^ which primitive cells were initially lacking. How protocells could have attained over time the ability to expediently perform communication and division, is a pending question.

Recently, we reported spontaneous formation of protocell-nanotube networks following a set of autonomous shape transformations on solid substrates^10^. The resulting structure is a population of surface-adhered protocells interconnected with lipid nanotubes. The nanotubular structures within the networks resemble the tunneling nanotubes (TNTs) between mammalian cells, which enable direct communication by transporting signaling molecules and even organelles^11, 12^. TNTs are also observed in bacterial cells, and likely provide an alternative route of intercellular exchange of cytoplasmic molecules and plasmids^13-15^. Whether the nanotubes in protocell networks^10^ would allow the transport of molecules, e.g. via molecular diffusion^16^, was initially not established.

In this work, we have investigated the ability of lipid nanotubes in surface-supported networks to transport prebiologically relevant constituents between the compartments: small water-soluble molecules, RNA and DNA. The protocell networks were formed for that purpose from label-free lipid membranes, allowing the focus of observations to be solely on fluorescently labeled cargo. Our findings confirm that the nanotubes function as tunneling interconnections between protocells, and are capable of transporting molecules and genetic polymers diffusively among them. We also characterized, supported by a dynamic analytical model, key physical parameters that are influential for the transport process. We hypothesize that nanotubes could have established a feasible means of communication and replication between prebiotic protocells, as in the investigated environment identical RNA and DNA fragments are easily distributed to nearby network nodes.

## Result and Discussion

### Formation of protocell-nanotube networks free of fluorophore

We started our experiments by preparing protocell-nanotube networks (PNNs) according to the protocol by Köksal et al.^10^. A schematic drawing and a fluorescence micrograph of a fluorescently labeled PNN (control) is shown in **Fig. 1a** and **Fig. 1b** respectively. Thereafter we prepared PNNs from label-free lipid membranes in order to exclusively visualize cargo molecules in the network without crosstalk^17^ with membrane fluorescence.

**Figure 1.**
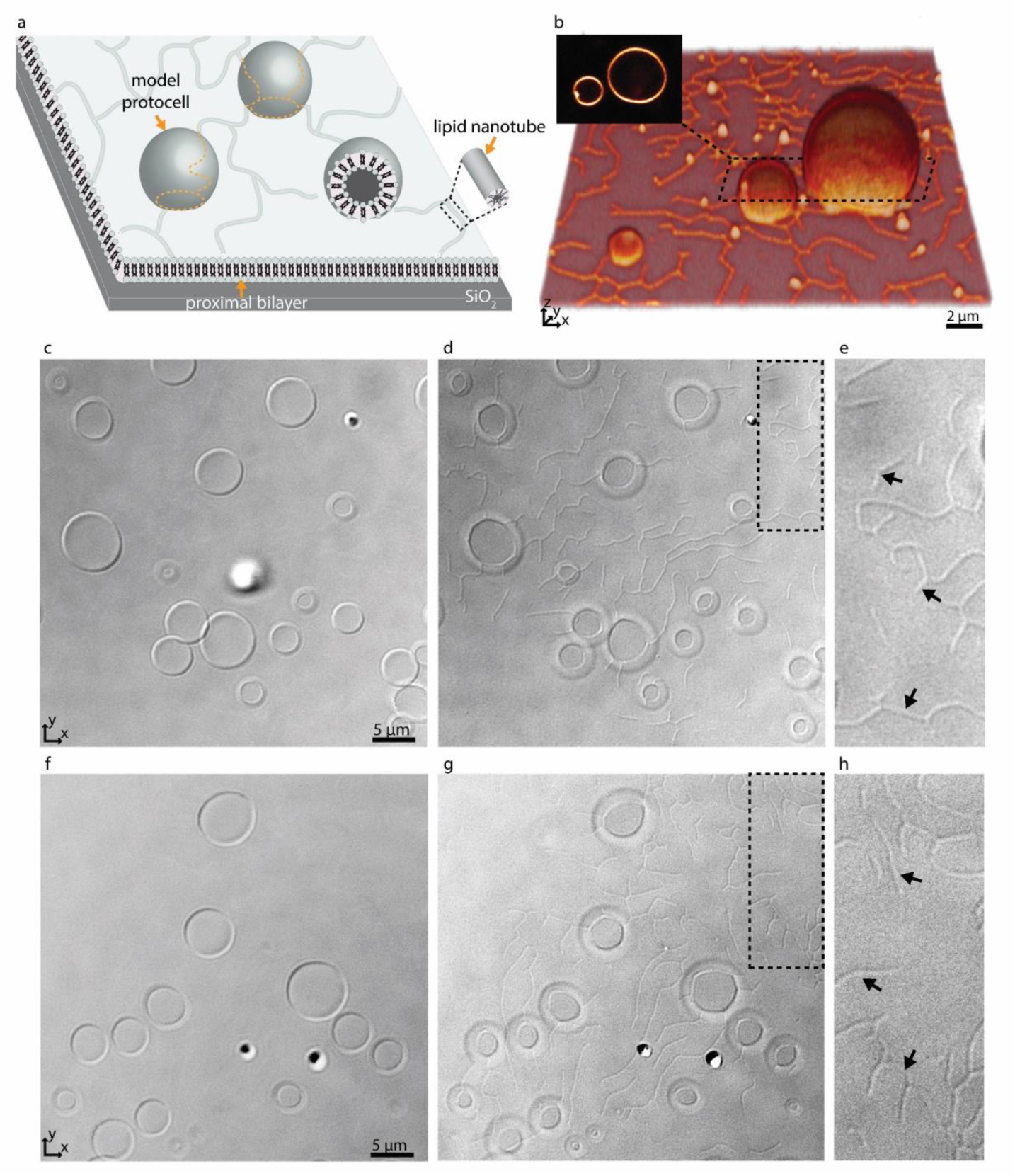
Protocell-nanotube networks (PNNs). (**a**) Schematic drawing of a protocell-nanotube network. The network and compartments are adhered to a surface supported bilayer. (**b**) 3D reconstructed confocal fluorescence micrograph of a PNN formed from a fluorescently labeled membrane (control). The inset shows a xy cross section of the two adjacent vesicles. (**c-h**) Differential interface contrast (DIC) microscopy images showing PNNs formed from unlabeled lipid membranes. DIC micrographs showing sections from the equator (**c, f**), and base (**d, g**) of the PNNs. The magnified version of the regions framed in dashed lines in (**d, g**) are shown in (**e, h**), revealing several nanotubes (black arrows).

We used differential interference contrast (DIC) microscopy to visualize the label-free PNNs (**Fig. 1c-h**), and the lipid nanotubes therein (**Fig. 1d-e, g-h**). The lack of fluorophore-conjugated lipids appeared to not interfere with the spontaneous formation of phospholipid protocell-nanotube networks. Overall, the findings were consistent with the PNNs formed from lipid preparation containing labeled phospholipids (**Fig. 1b**).

### Encapsulation and transport of cargo

Following the formation of PNNs, we introduced fluorescent cargo to the ambient solution in the vicinity of the compartments by means of an open-volume microfluidic device^18, 19^. A selected sample region was locally superfused with an aqueous buffer containing the cargo-molecules (**Fig. 2a)**. The supplied cargo molecules were ATTO 488 (**Fig. 2a**), a 10-base RNA labeled with fluorescein amidite (FAM), or a 20-base single stranded DNA, also labeled with FAM. During superfusion, these molecules passed the membrane and entered the protocell-nanotube networks through transient nano-pores^10, 20^. The concentration of FAM-RNA and FAM-ssDNA inside the compartments after 4 min. of superfusion was observed to be lower compared to ATTO 488 (**Fig. S1**). This is likely due to the higher molecular weight of the RNA and DNA, slowing down their diffusion through the pores (6.8 kDa for FAM-ssDNA, 3.4 kDa for FAM-RNA vs. 0.8 kDa ATTO 488).

**Figure 2.**
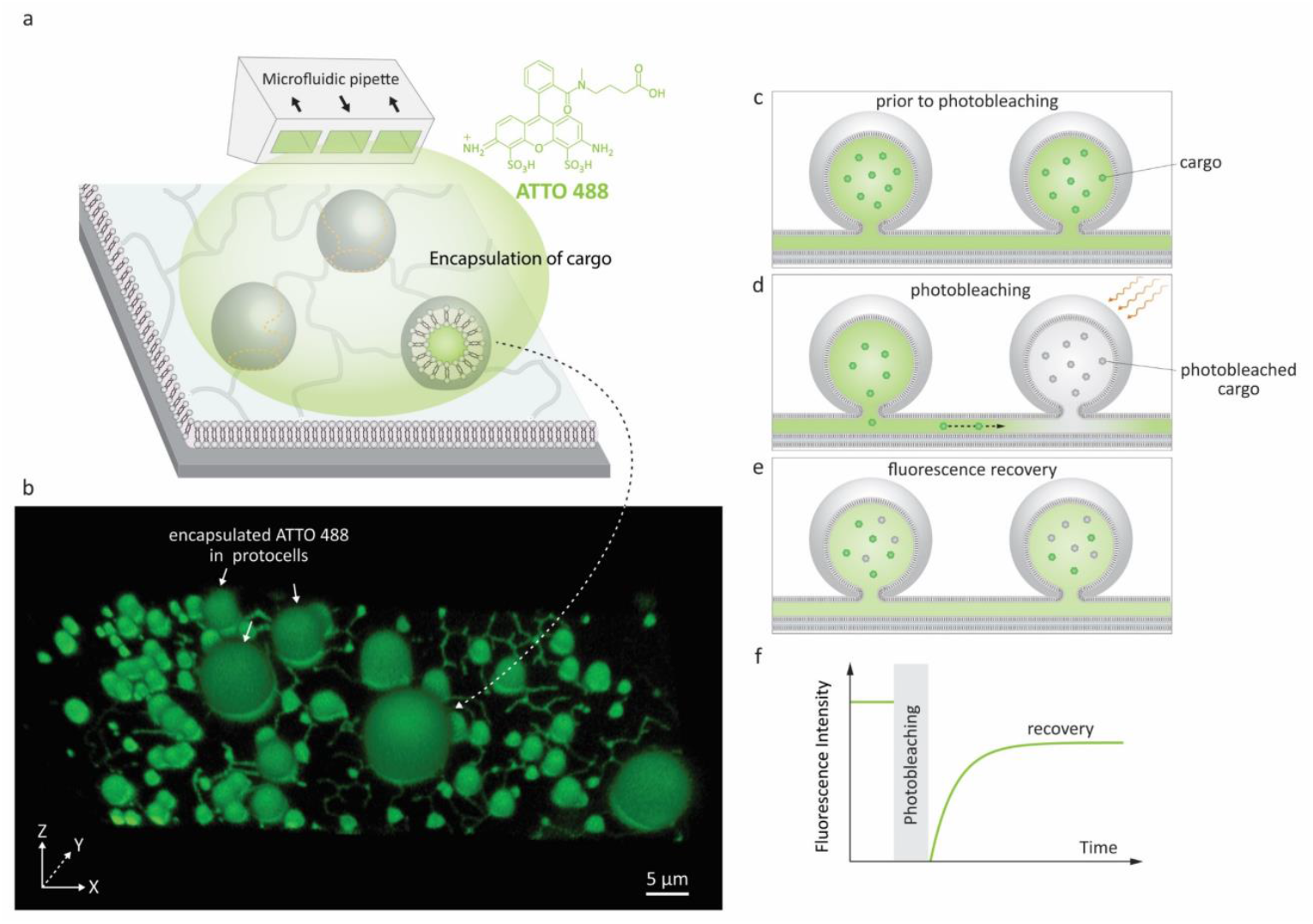
Encapsulation of molecular cargo inside PNN and FRAP experiment. (**a**) Schematic representation of the encapsulation experiment. An open microfluidic device (microfluidic pipette) creates a confined exposure zone around the protocells, delivering different cargo molecules, e.g. fluorescent dye ATTO 488. (**b**) 3D confocal micrograph of a label-free PNN upon encapsulating fluorescent dye inside the nanotubes and the protocells. (**c-e**) Schematic drawing of the FRAP (fluorescence recovery after photobleaching) experiment. (**c**) The fluorescent cargo is encapsulated inside the protocells and the connecting nanotubes, corresponding to **(b**). (**d**) Cargo in one of the protocells in **(a)** is photobleached using high laser intensity. (**e**) The fluorescence recovers due to the diffusion of fluorescent cargo from neighboring protocell through the nanotube. (**f**) Schematic graph depicting typical FRAP curve.

After ∼4 min of superfusion, the encapsulated cargo could be observed within the model protocells and nanotubes of the network via confocal microscopy. The confocal micrograph presented in **Fig. 2b** shows encapsulated ATTO 488 inside a surface-supported PNN. It confirms that the cargo molecules can enter the nanotubes, but cannot alone affirm their ability to transport the cargo. There remains still the possibility that the nanotubes are not conducting, but contain membrane defects which could block the tubes and prevent the exchange of molecules between the compartments.

After encapsulating the cargo, we investigated the transport of the molecules within the network by means of fluorescence recovery after photobleaching (FRAP) experiments (**Fig. 2c-f**). FRAP is a commonly used technique in cell biology and biomaterials science to determine dynamic processes e.g. membrane fluidity^12^, protein localization and mobility^21^, protein trafficking in intercellular nanotubes^22, 23^ and intracellular protein transport in ER or Golgi^24, 25^. When an isolated lipid vesicle encapsulating a fluorescent solution suspended in a non-fluorescent aqueous environment is photobleached, its intensity does not recover as there is no access to a new source of fluorophores for replenishment (**Fig. S2**). If the vesicles are physically connected through the tunneling nanotubes (**Fig. 2c-d**), and the molecules are able to diffuse through the tubes, the fluorescence signal of the photobleached vesicles recovers (**Fig. 2e-f**).

We verified this hypothesis by encapsulating ATTO 488, FAM-RNA and FAM-ssDNA in several nodes of the protocell networks, photobleaching selected compartments in the network and subsequently measuring the recovery of the initially photobleached compartments. The results from multiple experiments are shown in **Fig. 3**. Each plot in graphs **Fig. 3 a, h, l**, labeled with a capital letter, shows the fluorescence recovery of a single compartment within a network. **Fig. 3a-g** are associated with the experiments using ATTO 488, **Fig. 3h-k** RNA, and **Fig. 3l-o** DNA as encapsulated molecular cargo. **Fig. 3b-d** and **e-g** show confocal microscopy time series from two different experiments. For each experiment, the recovery of the compartments encircled in dashed lines was monitored and plotted in **Fig. 3a** (J and B). Plot E in **Fig. 3h** is obtained from the lipid compartment shown in **Fig. 3i-k**, and plot C in **Fig. 3l** from the compartment in **Fig. 3m-o**. Confocal microscopy time series corresponding to all other plots shown in **Fig. 3** are presented in **Fig. S3-5**.

**Figure 3.**
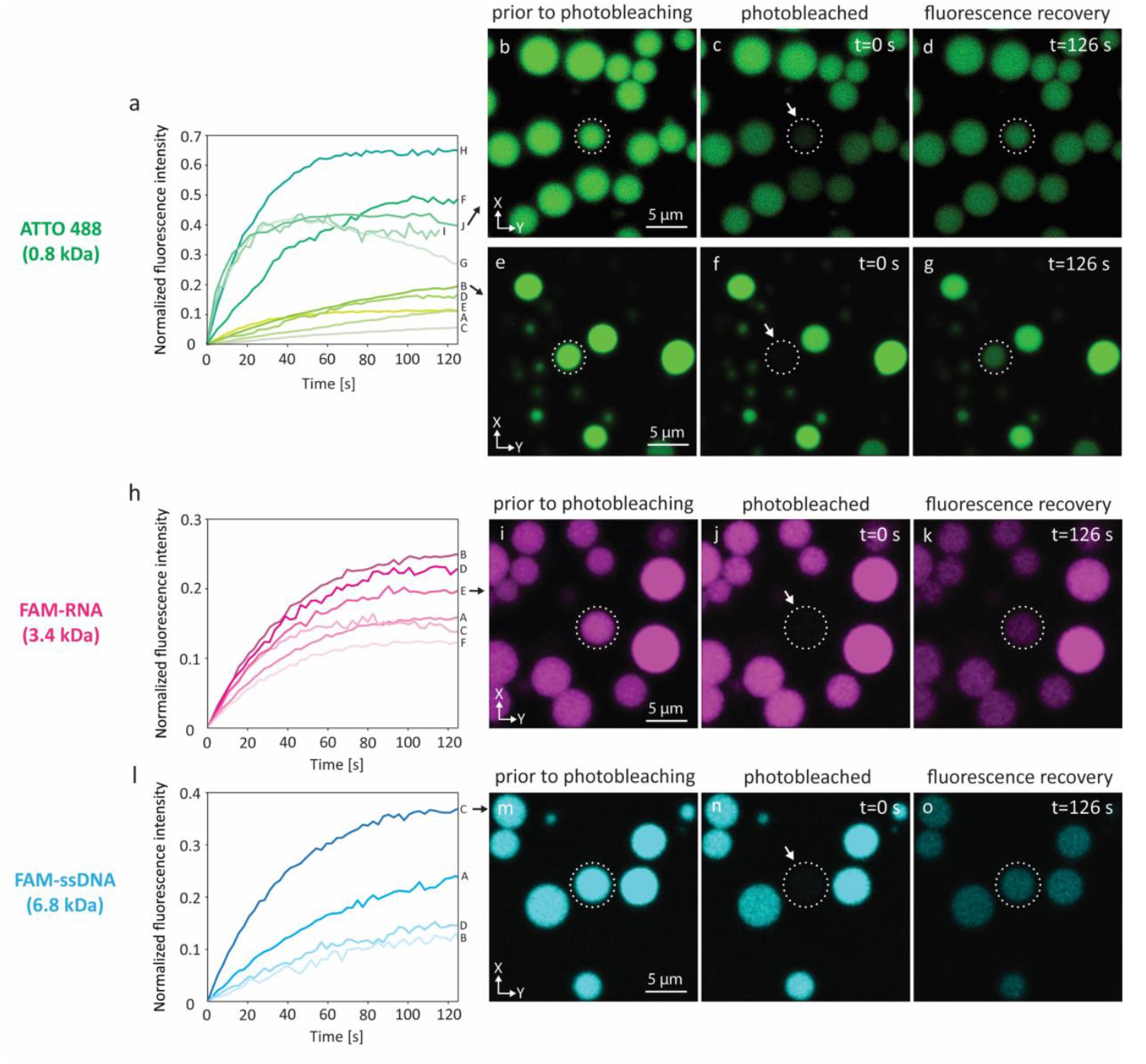
FRAP of selected compartments within PNNs. Three different cargo molecules, ATTO 488 (green color), FAM-RNA (magenta color) and FAM-ssDNA (cyan color) were loaded into protocell networks. (**a**) Plots show 10 FRAP experiments of ATTO 488 (A-J), each plot representing one experiment. (**b-d**) Confocal micrographs corresponding to plot J in panel (a): **(b)** ATTO 488-containing model protocell prior to, **(c)** during, **(d)** after, photobleaching. (**e-g**) Micrographs of the experiment corresponding to plot B in panel (a). (**h**) FRAP curves for RNA (plots A to F). (**i-k**) Micrographs of the FRAP experiment corresponding to plot E in panel (h). (**l**) Plots showing 4 FRAP experiments for DNA (plots A to D). Micrographs of experiment C are shown in (**m-o**). The compartments monitored for recovery are encircled in white dashed lines. All plots are normalized to the fluorescence intensity prior to the photobleaching.

Plots A-E in **Fig. 3a** show a final recovery of ∼5-20% of the initial fluorescence intensity. Plots F-J show more rapid recovery up to 65% (plot H) of initial intensity. The compartment represented in plot I could only be monitored until 115 s and further data collection could not be achieved. Plots G and J show a decline after reaching 40% of the initial intensity. Decrease in fluorescence intensity can be due to inherent photobleaching caused by continuous imaging^26^, or leakage from the compartments into the ambient solution via transient pores or defects in the membrane^27, 28^. Another possibility is transport of content from the recovered vesicle to adjacent compartments through other established nanotubular connections. It is challenging to determine to which exact compartments transfer of material would occur, as protocell-nanotube networks are extending out of the field of view for tens to hundreds of micrometers.

RNA containing compartments (**Fig. 3h-k, Fig. S4**) recover ∼12–26% of the original intensity over a time period of 100–120 s. It was more challenging to internalize DNA inside the network compared to ATTO 488 and RNA, due to their comparatively larger size. We conducted a total of four FRAP experiments with DNA (**Fig. 3l-o, Fig. S5**). The amount of recovery varied between ∼12% and 36%. We observed the transfer of all cargo molecules between the compartments, and established thus proof of principle.

### Geometric parameters influencing transport

We employed a simple analytical model to determine the impact of certain geometrical parameters for the molecular transport within the protocell-nanotube networks (**Fig. 4**). Although other transport mechanisms between vesicles via nanotubes have been reported, e.g. Marangoni transport^29^, for the model we mainly focus on molecular diffusion as means of transport. Rate equations describing the equilibration of particles between two^30, 31^ or more^31^ chambers connected with a capillary (**Fig. 4a)** were previously established. Volume of the compartments, diffusion coefficient of the molecules traveling between the compartments, length and radius of the connective nanoconduits were taken into account to calculate the diffusion rate^30, 31^ and relaxation time^30^.

**Figure 4.**
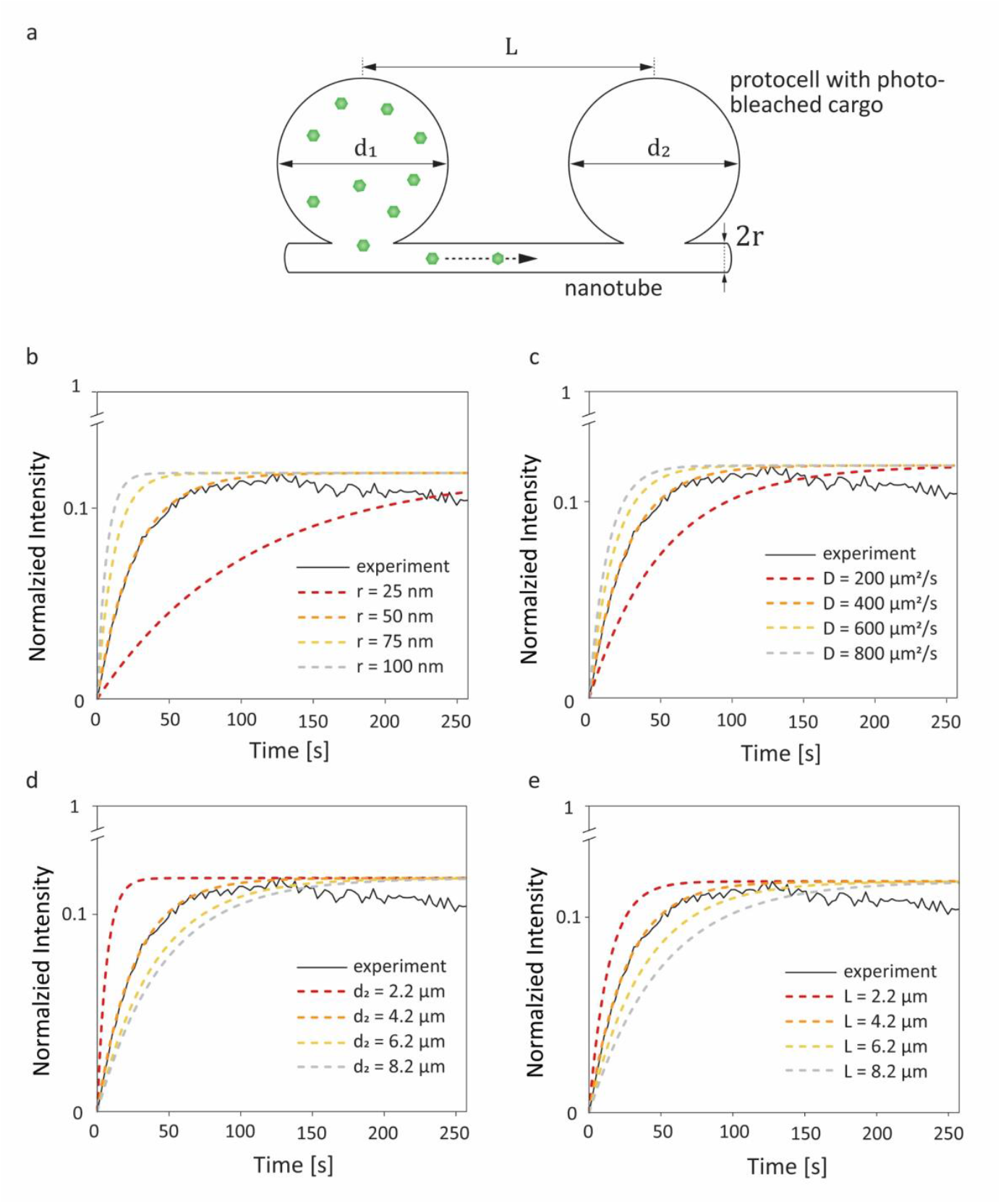
Geometric parameters influencing molecular diffusion in a two-vesicle system. (**a**) The analytical model is based upon two vesicles with diameters *d*_1_ and *d*_2_, connected through nanotube with a radius of *r* and distance *L*. ATTO 488 transports to the photobleached vesicle. Plot depicted in a black continuous line is experiment shown in **Fig. S6**. Dashed lines are obtained with different variables applied to Eq. 3. Curve fits to the recovery of the photobleached vesicle (solid line) while changing the (**b**) tube diameter, (**c**) diffusion coefficient of the cargo molecule, (**d**) compartment size, and (**e**) nanotube length. The plots are normalized to the fluorescence intensity prior to the photobleaching.

The above-mentioned models are based on simple, closed systems (dead ends) containing straight tubular connections^30-32^ (**Fig. 4a)**, where the networks in our experiments typically consists of multiple protocells with a branched, complex network of nanotubes extending outwards with many inter-connections and junctions (**Fig. 1**). Another feature that is different in our experiments is that the receiving (photobleached) lipid compartments are not free of cargo but contain the cargo with quenched fluorophores (**Fig. 2d**). Despite these differences, the models appear to provide a good approximation to the experimental system we have.

We compared the fluorescence recovery time of the compartment shown in **Fig. S6** to the relaxation time predicted by the model^30^ (**Fig. 4b-e**) (*cf*. Materials and Methods for details of the model). The two adjacent compartments in the experiment contain ATTO 488, which has a diffusion coefficient (*D*) of 400 μm^2^/s ^33^. We assume that the protocell compartments are spherical and connected with a straight tube of length *L* between the centers of their base. Each of the two compartments has a diameter of 4.2 μm and we take *L* = 4.2 μm since the compartments are located very close to each other **(Fig. S6)**. The radius of the nanotube, r, is the unknown variable.

Recovery time based on nanotubular connections with varying nanotube radii were plotted while keeping *d*_1_ = 4.2 μm, *d*_2_ = 4.2 μm, *D* = 400 μm^2^/s, *L* = 4.2 μm (*cf*. Eq. 1 and Eq. 3 in Materials and Methods) (**Fig. 4b**). The best match of the experimental curve to the theoretical curve was achieved for the tube with a radius of 50 nm (**Fig. 4b**). This is within the range of tunneling nanotubes observed in cellular^11^ and artificial systems^34^.

Next, we focused on diffusion coefficient of the cargo molecule. The expected relaxation/recovery times for varying *D* values are plotted in **Fig. 4c**. The best fit was obtained for *D* = 400 μm^2^/s, which is the diffusion coefficient of ATTO 488. For a 10-base RNA *D* = 200 μm^2^/s ^35^, and for a 20-base ssDNA *D* = 152 μm^2^/s ^36^. We ignored the contribution of the FAM label that is conjugated to the nucleotides. The recovery of RNA and DNA inside the compartments is expected to be slower than of ATTO 488; the recovery curve would be similar to the one shown with a red-dashed line in **Fig. 4c**, where *D* = 200 μm^2^/s.

**Fig. 4d** shows multiple plots corresponding to compartments with varying sizes. As anticipated, the larger the diameter of the compartments, the more time it takes to reach equilibrium. For the receiving (photobleached) vesicle this means more volume needs to be filled, while for the donating vesicle it means a lower probability for the cargo to reach the entrance to the nanotube.

Finally we investigated the length of the nanotube (**Fig. 4e**). With increasing nanotube length the travel time of the cargo in the nanotube increases, resulting in slower fluorescence recovery (**Fig. 4e**). The nanotube length, related to the distance between the protocells, is directly proportional to the relaxation time (**Eq. 3**). The exact length of the connecting nanotubes in the experiments is difficult to predict, as they are almost never straight due to pinning and branching^37, 38^.

The recovery curve of the experiment shown in **Fig. S6** (plot shown with a continuous black line in **Fig. 4b-e**) declines gradually over time. This can be due to inherent photobleaching caused by continuous imaging^26^, or leakage from the compartments^27, 28^ as discussed above.

## Conclusion

Our findings confirm that the lipid nanotubes in surface-adhered protocell networks are open, and allow molecular transport between the interconnected bilayer-encapsulated compartments. The rate of the diffusive transport is highly dependent on the structure of the network and influenced by the parameters such as the radius and length of the nanotubes, size of the compartments and the diffusion coefficient of the molecule that is transported. There is a physical limit to how small the radius of the lipid nanotubes can be without the presence of curvature stabilizing proteins, rendering the other parameters deciding factors for the diffusion rate in a prebiotically relevant context. We conclude that it appears feasible to increase complexity by encapsulating reactants for prebiotic reactions in PNNs, in order to gain a deeper understanding of possible chemical communication processes within primitive cell populations at the origin of life.

## Materials and Methods

### Lipid preparation

Lipid suspensions were prepared with soybean polar extract and E. coli polar extract (Avanti Polar Lipids, USA) (50:50 wt %), using the dehydration-rehydration method^39^. Briefly, lipids were dissolved in chloroform in a 10 mL pear shaped bottom flask leading to a final concentration 10 mg/mL. For the fluorescently labeled sample shown in **Fig. 1b**, 1 wt % of lipid conjugated fluorophore 16:0 Rhod Liss PE (Avanti Polar Lipids, USA) was added into the lipid mixture. 300 μL of the dissolved lipid mixture was placed in a rotary evaporator and the solvent was removed at 24 rpm and reduced pressure (20 kPa) for 6 hours to form a dry lipid film. The dry lipid film was rehydrated with 3 mL phosphate-buffered saline (PBS) followed by addition of 30 μL glycerol. The PBS contained 5 mM Trizma base, 30 mM K_3_PO_4_, 30 mM KH_2_PO_4_, 3 mM MgSO_4_·7H_2_O, and 0.5 mM Na_2_-EDTA (pH = 7.4, adjusted with H_3_PO_4_). The lipid suspension was kept at 4 °C overnight to allow swelling of the lipid cake. The following day, the suspension was sonicated for 5–10 s at room temperature, leading to formation of giant vesicle suspension. The suspension was aliquoted and stored at -18 °C.

### Surface preparation

SiO_2_ surfaces were fabricated at the Norwegian Micro- and Nano-Fabrication Facility at the University of Oslo (MiNaLab). Thin films were deposited onto glass cover slips (Menzel Gläss #1, 100-150 μm thickness; WillCo Wells B.V., Amsterdam, NL). SiO_2_ films were deposited by E-beam physical vapor deposition using an EvoVac instrument (Ångstrom Engineering, Canada), to a final thicknesses 84 nm.

### Sample preparation

For sample preparation, two 4 μL droplets of lipid suspension were placed on a clean glass and dehydrated in desiccator for 20-25 min under low pressure. The dry lipid film was rehydrated for 10 min with 0.5-1 mL of Na-HEPES buffer containing 10 mM HEPES and 100 mM NaCl (pH = 7.8, adjusted with NaOH). The rehydrated suspension was later transferred into observational chamber containing Ca-HEPES buffer (10 mM HEPES, 100 mM NaCl and 4 mM CaCl_2_, pH = 7.8, adjusted with NaOH). The sample was kept at room temperature for 2-3 days for PNN formation and protocell growth. Alternatively, to speed up the protocell growth, the sample was incubated in 35°C until the following day to promote protocell growth^38^.

### Encapsulation of cargo molecules

The cargo molecules were delivered with an open volume microfluidic pipette^18, 19^ (Fluicell AB, Sweden) positioned using 3-axis hydraulic micromanipulator (Narishige, Japan) to the vicinity of the protocell-nanotube structures. The protocells were superfused with solutions of Ca-HEPES buffer containing 500 μM of ATTO 488 carboxyl (Atto-Tech GmbH, Germany). DNA solution was prepared and delivered in nuclease-free water (Thermo Fisher Scientific, USA) by dissolving 200 μM of 20-base ssDNA (5’/56-FAM/TGT ACG TCA CAA CTA CCC CC-3’, Integrated DNA Technologies, USA). 10-base RNA oligomers (5′-FAM-AAA AAA AAA A-3′, Dharmacon, USA) were dissolved in nuclease-free water to a final concentration of 100 μM. Exposing the nanotube networks rapidly to nuclease-free deionized water -a hypotonic environment-facilitated the rapid swelling and growth of membranous compartments (**Movie S1**), reducing the time period of growth from hours^10^ to minutes.

### FRAP curve fitting

The theoretical FRAP curves in **Fig. 4, Fig. S6** were obtained using MATLAB R2020b. The fluorescence recovery, *F*, over time, *t*, takes the form of an exponential function (**Fig. 2f**) and the FRAP curve can be fitted to^40^:

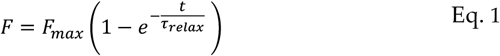

where *F*_max_ is the maximum fluorescence recovery and *τ*_relax_ is the relaxation time, which is the time for the system to reach chemical equilibrium (*F*_max_) at constant, initial diffusion rate. In reality the diffusion rate decreases over time as the concentration gradient between two compartments decreases. The relaxation time (Eq. 1) stands for the duration of time at the end of which, the fluorescence recovery reaches (1-1/e) of *F*_max_.

For a simple system with two spherical vesicles, the relaxation time, *τ*_relax_, is given by^30-32^:

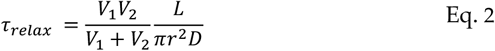

where *V*_1_ and *V*_2_ are the volumes of the compartments between which the diffusion takes place. The diffusion coefficient, *D*, is distinct for each molecule, while *r* and *L* are the radius and length of the nanotube. Because *V* = *πd*^3^/6, this expression can be written using the diameters, *d*, of the vesicles:

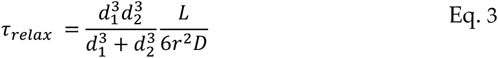

### Microscopy imaging and analysis

Imaging has been performed using Laser scanning confocal microscope DMi8 (Leica Microsystems, Germany). The 3D fluorescence micrograph in **Fig. 1** was reconstructed using the Leica Application Suite X Software (Leica Microsystems, Germany). Image enhancement of fluorescence micrographs for the figures was performed with the Adobe Photoshop CS4 (Adobe Systems, USA). All FRAP curves in **Fig. 3, 4, S2, S6** were plotted in MATLAB R2020b. Schematic drawings were created with Adobe Illustrator CS4 (Adobe Systems, USA).

## Acknowledgements

This work was made possible through financial support obtained from the Research Council of Norway (Forskningsrådet) Project Grant 274433, UiO: Life Sciences Convergence Environment, as well as the startup funding provided by the Centre for Molecular Medicine Norway (RCN 187615), and the Faculty of Mathematics and Natural Sciences at the University of Oslo.

