## Supporting Information for "Transport among protocells via tunneling nanotubes"

### S1. Encapsulation of cargo molecules

Three different fluorescent cargo molecules with varying molecular weights were introduced to the protocell-nanotube networks (PNNs) using an open-space microfluidic pipette[1, 2]: a fluorescent dye (ATTO 488) (**Fig. S1a-b**), RNA (**Fig. S1c-d**) and DNA (**Fig. S1e-f**). Panels **a**,**c** and **e**, represent the part of the experiment during which the fluorescently labeled cargo molecules are continuously exposed to a region on the PNNs. Panels **b**,**d** and **f** shows the networks after ~4 min of exposure.

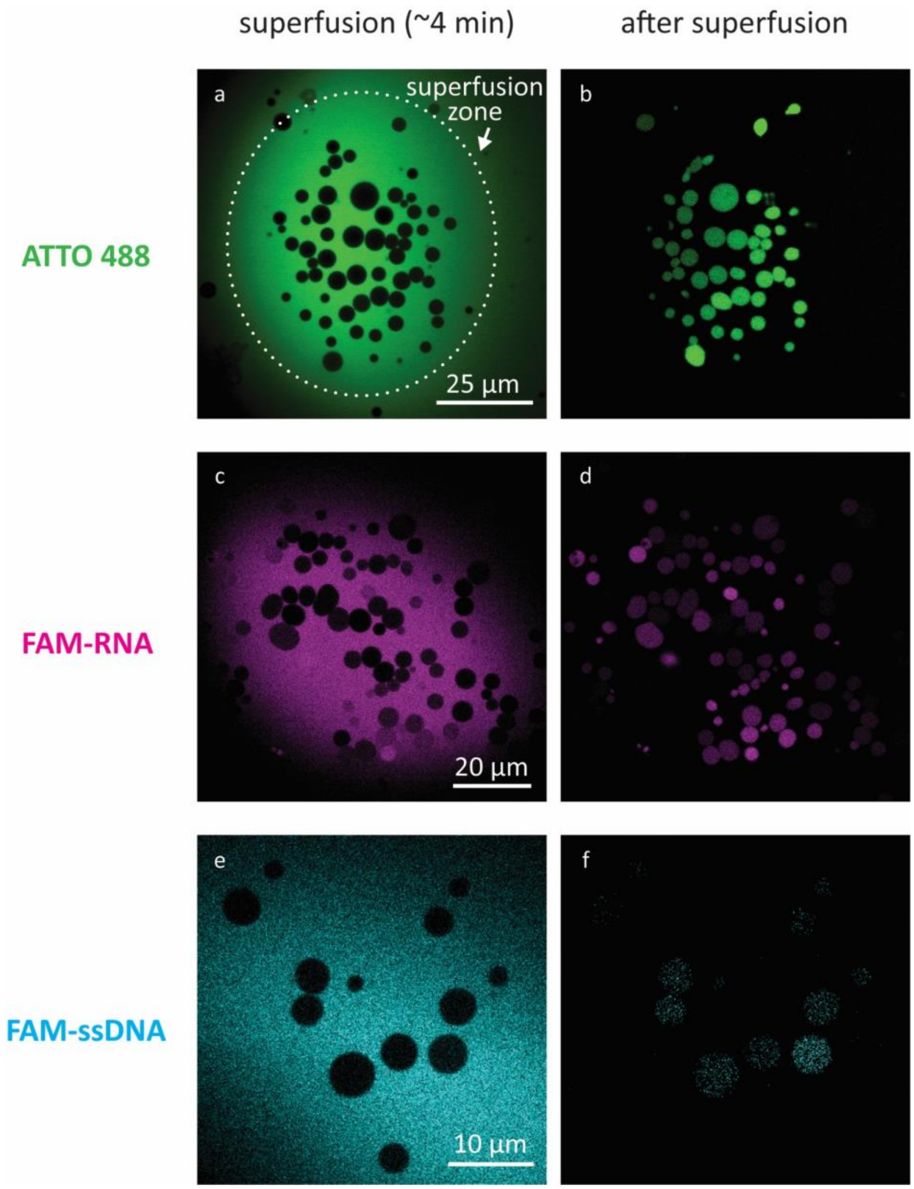

**Figure S1.** Encapsulation of different cargo molecules. **a, c, e**) Confocal micrographs ~4 min into the superfusion, and **b, d, f**) right after superfusion is terminated. (**a-b**) ATTO 488, (**c-d**) FAM-RNA, (**e-f**) FAMss-DNA.

### S2. FRAP of an isolated vesicle (control)

We performed a control FRAP experiment on an isolated, surface-adhered giant unilamellar vesicle (GUV) containing ATTO 488 (**Fig. S3a-b**). Upon photobleaching, no recovery was observed (**Fig. S3c-d**). This result confirms necessity of a nanotubular connection for recovery of the fluorescence intensity of a lipid compartment in PNNs.

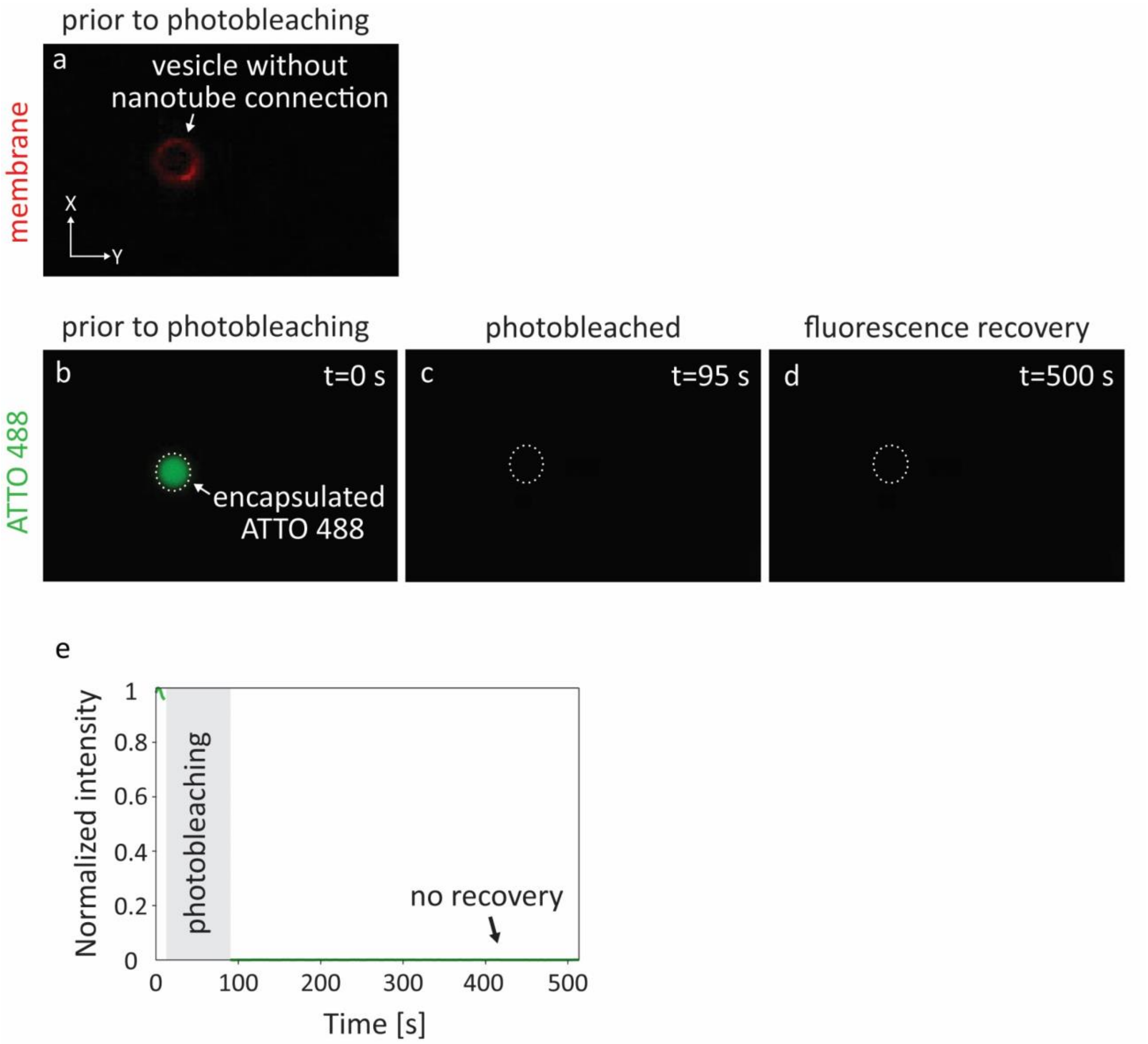

**Figure S2.** FRAP of an isolated GUV on a solid substrate. (**a-b**) Confocal micrograph of an isolated GUV, encapsulating ATTO 488. (a) shows the membrane fluorescence, and (b) the fluorescence of the internalized dye, ATTO 488. Photobleached GUV (**c-d**). (**e**) FRAP curve corresponding to (**b-d**). The plot is normalized to the fluorescence intensity prior to photobleaching. The diameter of the vesicle is 4 μm.

### S3. FRAP experiments

Confocal microscopy time series corresponding to the plots shown in **Fig. 3** of the main manuscript. Several FRAP experiments were performed for each cargo molecule ATTO 488 (**Fig. S3**), RNA (**Fig. S4**) and DNA (**Fig. S5**). Each experiment is labeled with the capital letters matching the labels of the plots in **Fig. 3a,h,l**.

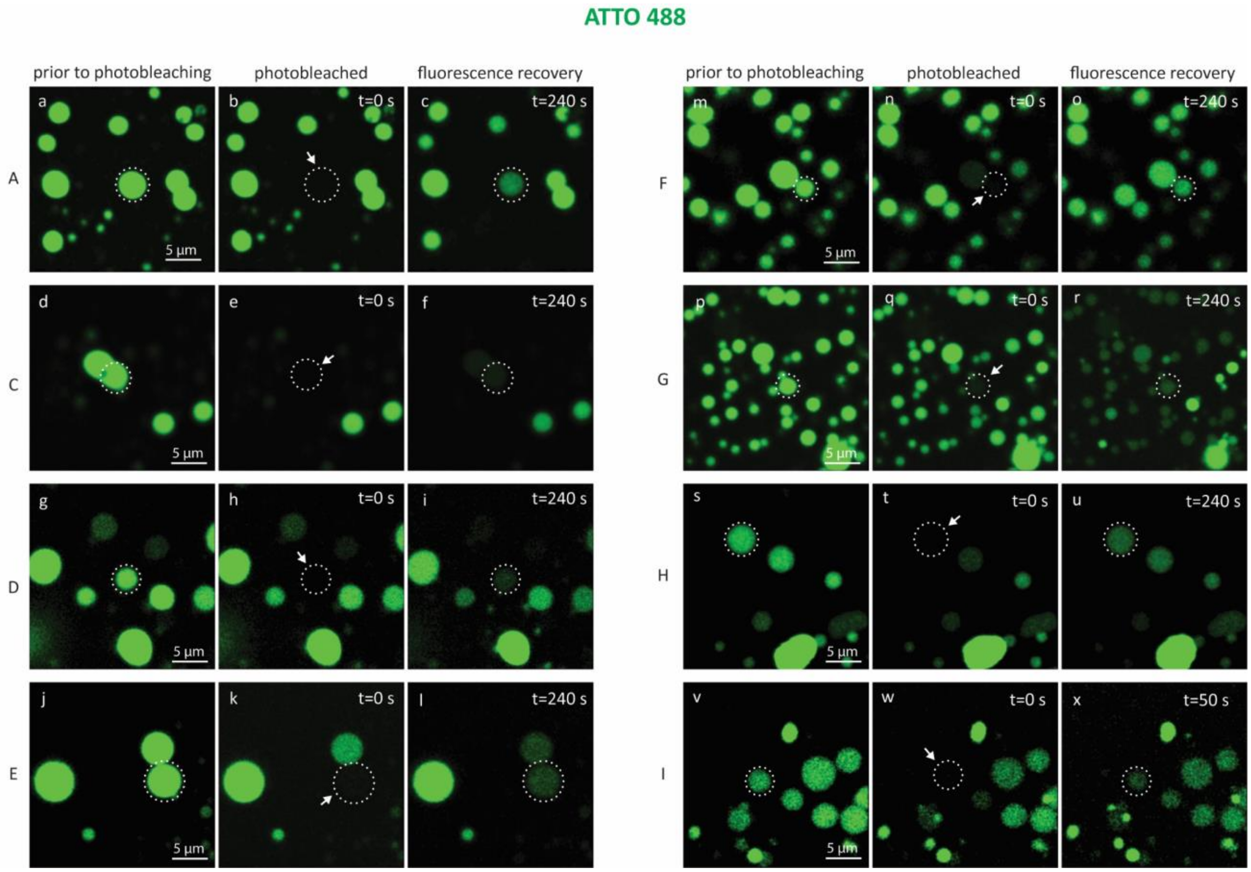

**Figure S3.** Confocal micrographs showing before, during and after photobleaching of compartments encapsulating **ATTO 488** in different experiments: A (**a-c**), C – I (**d-x**). Each set of micrographs show a model protocell targeted for photobleaching (encircled in dotted lines). Three time points in each experiment represent: prior to photobleaching, during photobleaching (arrows) and during fluorescence recovery.

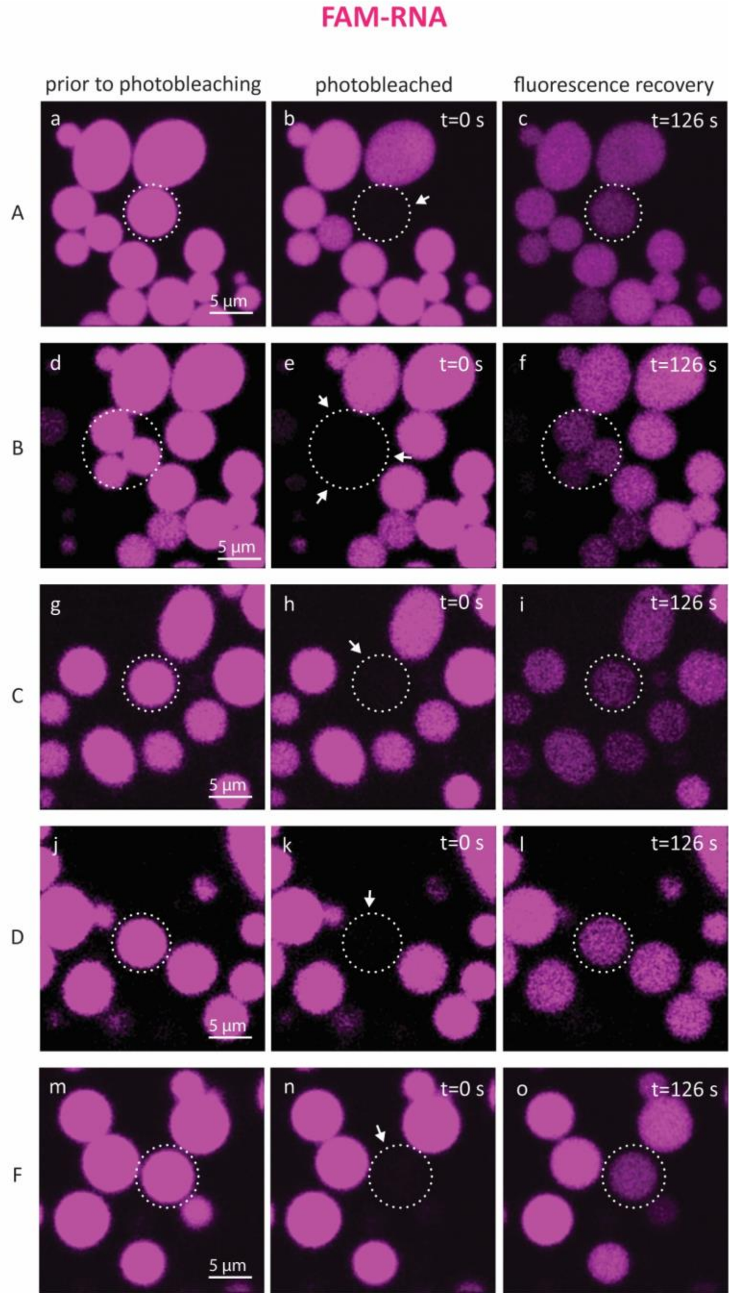

**Figure S4.** Confocal micrographs showing before, during and after photobleaching of compartments encapsulating **RNA** in different experiments: A-F (**a-o**). Each set of micrographs show a model protocell targeted for photobleaching (encircled in dotted lines). Three time points in each experiment represent: prior to photobleaching, during photobleaching (arrows) and during fluorescence recovery.

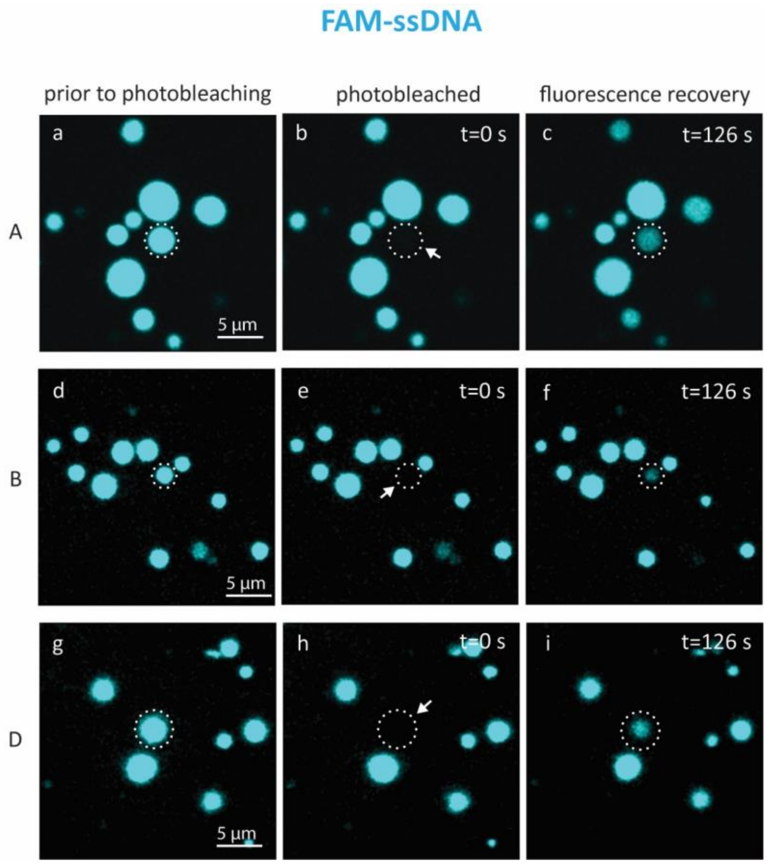

**Figure S5.** Confocal micrographs showing before, during and after photobleaching of compartments encapsulating **DNA** in different experiments: A-D (**a-i**). Each set of micrographs show a model protocell targeted for photobleaching (encircled in dotted lines). Three time points in each experiment represent: prior to photobleaching, during photobleaching (arrows) and during fluorescence recovery.

### S4. Fluorescence recovery in a two-compartment system

A FRAP experiment followed by the transport of ATTO 488 between two adjacent protocells has been presented in **Fig. S6a-c**. **Fig. S6d** shows the fluorescence intensity of the donor (yellow plot) and acceptor (green plot) protocell, over time. The dashed line is the theoretical fit based on a two-compartment model[3] (**Fig. 4**), which overlaps with the fluorescence recovery (green plot). Fluorescence intensity of both compartments after the recovery is below 50%, which is comparable to the findings of other numerical methods that describe the diffusive transport between two compartments[4]. The high fluorescence intensity of the leftmost protocell in **Fig. S6c** maintains during several minutes, indicating that it has no open nanotubular connection to protocell 1 or 2, and is not a contributing donor compartment (**Fig. S6a**).

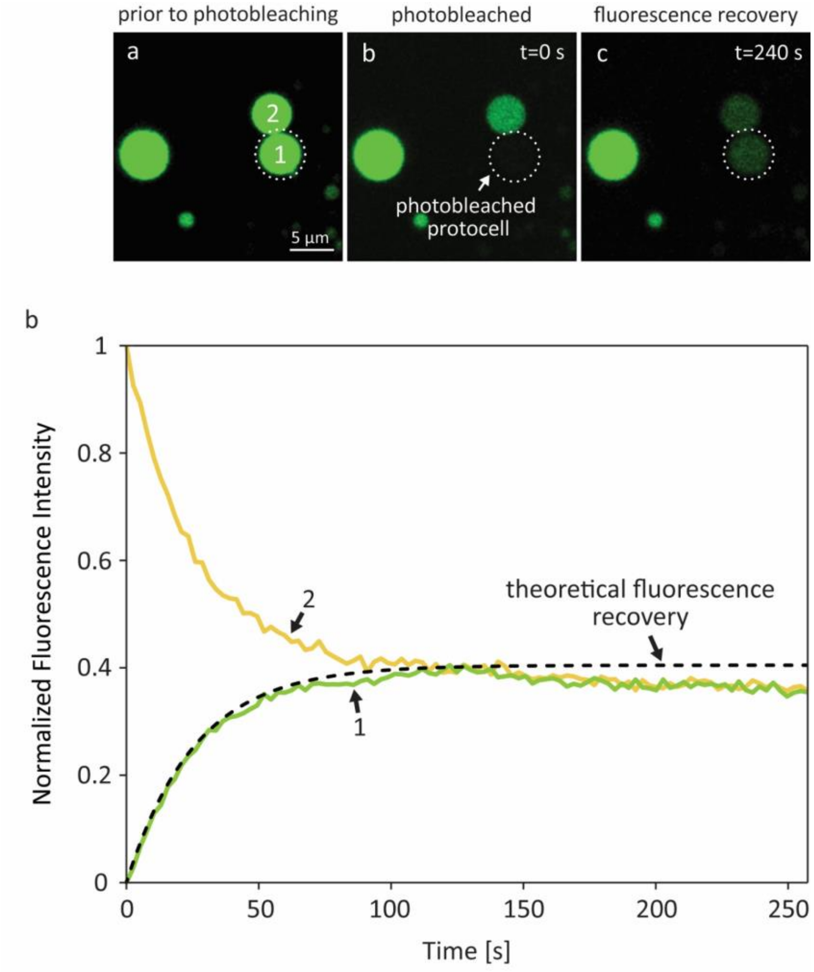

**Figure S6.** Fluorescence intensity of a two-compartment system after photobleaching of one of the compartments. (**a-c**) Protocell 1 (encircled in dotted line) is photobleached. (**d**) Fluorescence intensity of the donor (yellow plot) and acceptor (green plot) vesicle, over time. The plots are normalized to the maximum fluorescence intensity of the nearby protocell, and the minimum fluorescence intensity of the photobleached protocell during the fluorescence recovery period.

#### S5. Supporting Movie

**Movie S1. Rapid formation of protocells during DNA exposure.** Laser scanning confocal microscopy time series showing rapid formation and growth of protocells from the nanotube network during DNA exposure.
